# The double-layered structure of amyloid-β assemblage on GM1-containing membranes catalytically promotes fibrillization

**DOI:** 10.1101/2022.06.26.497640

**Authors:** Maho Yagi-Utsumi, Satoru G. Itoh, Hisashi Okumura, Katsuhiko Yanagisawa, Koichi Kato, Katsuyuki Nishimura

## Abstract

Alzheimer’s disease (AD) is associated with progressive accumulation of amyloid-β (Aβ) cross-β fibrils in the brain. Aβ species tightly associated with GM1 ganglioside, a glycosphingolipid abundant in neuronal membranes, promote amyloid fibril formation; therefore, they could be attractive clinical targets. However, the active conformational state of Aβ in GM1-containing lipid membranes is still unknown. The present solid-state nuclear magnetic resonance study revealed a nonfibrillar Aβ assemblage characterized by a double-layered antiparallel β-structure specifically formed on GM1 ganglioside clusters. Our data show that this unique assemblage was not transformed into fibrils on GM1-containing membranes, but could promote conversion of monomeric Aβ into fibrils, suggesting that a solvent-exposed hydrophobic layer provides a catalytic surface evoking Aβ fibril formation. Our findings will offer structural clues for designing drugs targeting catalytically active Aβ conformational species for the development of anti-AD therapeutics.

## Introduction

Neurodegenerative diseases, such as Alzheimer’s disease (AD), Parkinson’s disease, and Huntington’s disease, are ascribed to pathogenic molecular processes involving conformational transitions of amyloidogenic proteins into toxic aggregates such as oligomers and fibrils (*1-3*). In addition to certain oligomers being deleterious aggregates (*4, 5*), amyloid fibrils have been demonstrated to exert cytotoxicity and play key roles in the persistence, progression, and propagation of amyloid diseases (*6*). Cross-β structures have been found in amyloid fibrils from various amyloidogenic proteins, suggesting that in-register parallel β-sheet formation is a universal feature of amyloid fibril structures (*3*). Recent studies have shed light on the mechanism by which amyloid fibril surfaces can catalyze secondary nucleation (*7*). Self-catalyzed secondary nucleation is a monomer-dependent process whereby transient monomer binding to a fibril lateral surface accelerates aggregation, giving rise to cytotoxic oligomers and short fibrils. In some instances, lipid membranes can serve as platforms for conformational transition and subsequent fibril formation. Indeed, membranes can accelerate the rate of primary nucleation (*8*) as well as elongation of amyloid fibrils, leading to bilayer disruption by increased fibril load.

AD is associated with progressive accumulation of cross-β fibrils of amyloid-β (Aβ), mainly consisting of 40 or 42 amino acids, in the brain resulting in formation of extracellular senile plaques (*9, 10*). It has been reported that binding of Aβ fibrils to cell membranes inhibits long-term potentiation in mouse hippocampal brain slices (*11*). Yanagisawa et al. identified a unique Aβ species tightly associated with GM1 ganglioside, a glycosphingolipid abundant in neuronal membranes, in the cerebral cortices of human brains exhibiting early pathological AD changes (*12*). A monoclonal antibody, 4396C that specifically binds to GM1-bound Aβ species stained neurons in the cerebral cortices of AD brains (*13*). A recent mass spectrometry imaging study revealed that GM1 was present in the core region of amyloid plaques and that the amount of deposited Aβ correlated with that of GM1 in AD mouse models (*14*).

Various *in vivo* and *in vitro* studies have demonstrated that GM1-bound Aβ species act as endogenous seeds for cerebral Aβ fibril formation, accelerating Aβ assembly (*13, 15-18*). Furthermore, amyloid fibrils on GM1-containing liposomes were reported as more toxic than those formed in aqueous solution (*19, 20*). Interactions between Aβ and the GM1 cluster have thus far been characterized by nuclear magnetic resonance (NMR) spectroscopy and molecular dynamics (MD) simulation using model systems such as GM1-containing micelles, highlighting α-helix formation at the hydrophobic/hydrophilic interface (*21-28*). However, the active conformational state of Aβ in GM1-containing lipid membranes remains to be elucidated, as it has only been characterized as a conformational epitope recognized by the 4396C antibody (*13*). Intriguingly, this antibody has been shown to be cross-reactive with Aβ at the ends of growing fibrils (*13*); administration of its Fab fragments significantly reduced plaque formation in AD model mice (*29*), suggesting a common conformational epitope between Aβ molecules in GM1 clusters and those on the fibril ends.

Here, we report a structural characterization of GM1-bound states of Aβ molecules. We established a protocol for stabilizing active Aβ catalytic species reactive with the 4396C antibody, using a GM1-rich phospholipid membrane system. On this basis, we successfully unveiled the three-dimensional structures of those Aβ species by solid-state NMR spectroscopy together with MD simulation.

## Results

### Stabilization of active catalytic Aβ species

To characterize the structure of GM1-bound Aβ molecules, we first optimized the protocol to prepare solid-state NMR samples. An Aβ solution and GM1/1,2-dimyristoyl-sn-glycero-3-phosphocholine (DMPC) vesicle suspension were mixed on ice and immediately subjected to ultracentrifugation at 4°C to prevent amyloid fibril formation, thereby collecting only the vesicle-bound fraction as precipitate, which was immediately lyophilized (**Fig. 1A**). To confirm the absence of amyloid fibrils in the collected fraction, part of the lyophilizate of the Aβ/GM1/DMPC fraction was rehydrated and subjected to electron microscopy, which detected only GM1/DMPC vesicles but no amyloid fibrils (**Fig. 1B**). This was also confirmed by a thioflavin T (ThT) assay, indicating no fluorescence enhancement of rehydrated lyophilizates of Aβ/GM1/DMPC and Aβ/DMPC fractions even after prolonged incubation at 37°C (**Fig. 1C**). A free Aβ solution exhibited a gradual increase in ThT fluorescence intensity during incubation at 37°C after a lag time of about 24 h, but it showed a greater increase of ThT fluorescence in the presence of the rehydrated lyophilizate of the Aβ/GM1/DMPC fraction (**Fig. 1D**). These data indicated that the collected fraction contained no detectable free Aβ molecules and that the GM1-membrane-bound Aβ itself was not transformed into ThT-reactive fibrils but instead promoted amyloid fibrillization of monomeric Aβ. We examined the reactivities of the Aβ/GM1/DMPC fraction with specific monoclonal antibodies. In addition to 4396C, it reacted with 6E10 and 4G8, known to recognize the N-terminal segment (Asp1-Lys16) and the mid part (Leu17-Val24) of Aβ, respectively, but not with AB27, a C-terminal specific antibody. These data indicate that the N-terminal and middle segments of Aβ are exposed to the solvent while its C-terminal part is not accessible to the antibody (**Fig. 1E**). Furthermore, the rehydrated lyophilizate of Aβ/DMPC fraction did not enhance amyloid fibrillization of monomeric Aβ nor was it recognized by 4396C, indicating that only GM1-bound Aβ can act as seed for acceleration of fibril formation.

**Fig. 1.**
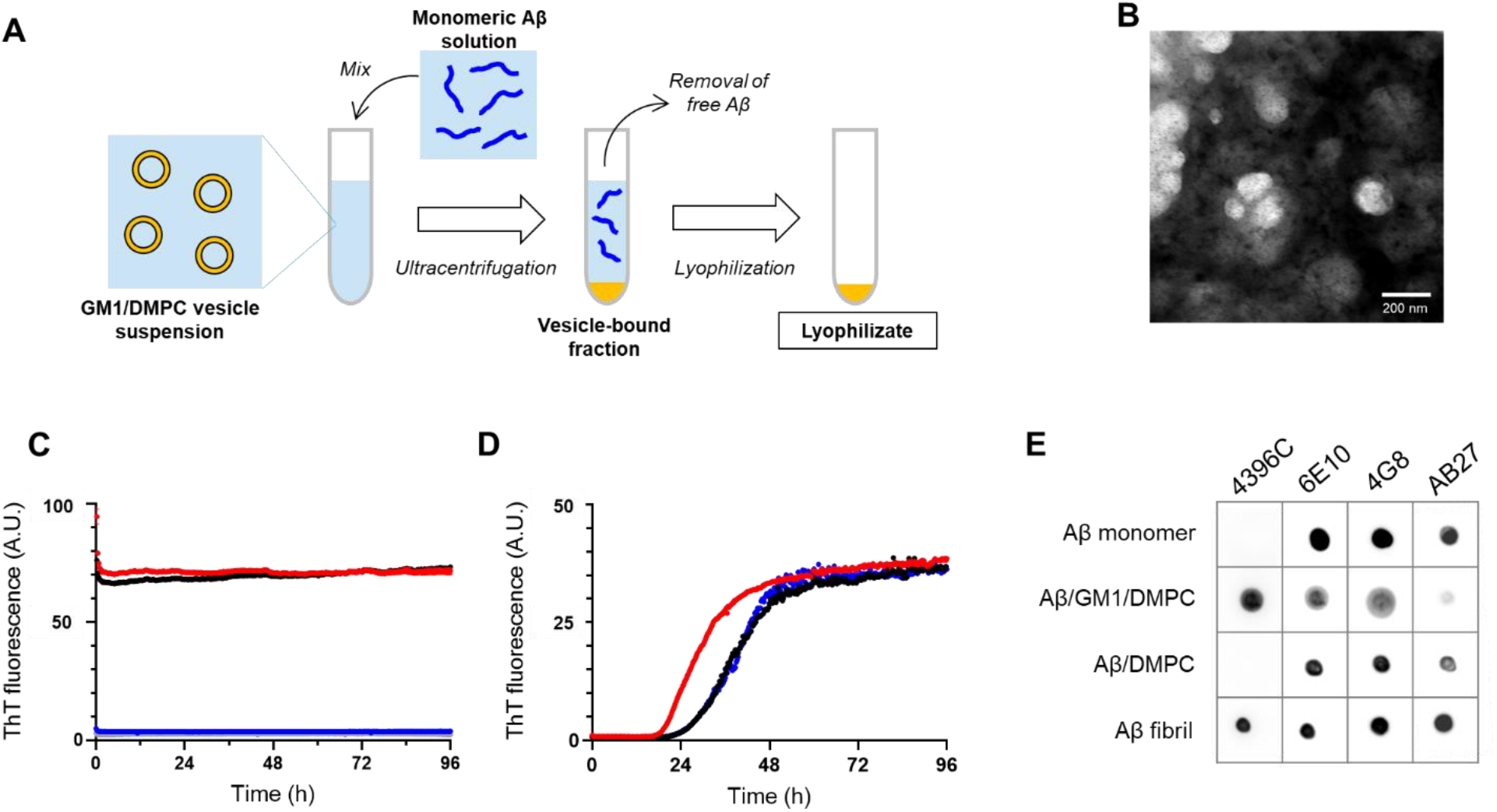
Preparation and characterization of active catalytic Aβ species. (**A**) Scheme of sample preparation of Aβ_1–40_ bound to GM1/DMPC vesicle for solid-state NMR. (**B**) TEM image of the Aβ_1– 40_/GM1/DMPC fraction. (**C**) ThT fluorescence assay of the rehydrated lyophilizate of Aβ_1– 40_/GM1/DMPC fraction (red), GM1/DMPC suspension (black), Aβ_1–40_/DMPC fraction (blue), and DMPC suspension (gray). (**D**) ThT fluorescence assay of Aβ_1–40_ in the absence (black) and presence of the rehydrated lyophilizate of Aβ_1–40_/GM1/DMPC fraction (red) or in the presence of the rehydrated lyophilizate of Aβ_1–40_/DMPC fraction (blue). (**E**) Dot blot assay of Aβ_1–40_ with monoclonal anti-Aβ antibodies. Monomeric Aβ_1–40_, the rehydrated lyophilizates of Aβ_1–40_/GM1/DMPC and Aβ_1– 40_/DMPC fractions, and preprepared Aβ_1–40_ fibril were blotted.

### Antiparallel β-structure of Aβ assemblage on GM1/DMPC vesicles

We characterized the conformation of isotope-labeled Aβ bound to GM1/DMPC vesicles based on solid-state NMR data. Sequential signal assignments were achieved by ^13^C dipolar-assisted rotational resonance/RF-assisted diffusion [DARR (*30*)/RAD (*31*)] experiment in conjunction with ^13^C-detected NCO and NCA heteronuclear correlation experiments using ^13^C**–**^15^N double cross polarization (DCP) (*32*) (**Fig. 2A, Figs. S1A and S1B**). In addition, the peaks originating from Val36 and Val40 were assigned by using site-specifically isotope-labeled Aβ (**Fig. S1C**). The results indicate that the observed peaks originated from Val12–Val40 of Aβ bound to GM1/DMPC vesicles (**Table S1**). No peak multiplicity was observed, indicating the conformational equivalence of this segment. Torsion angle prediction based on Aβ backbone chemical shifts revealed that the segments Leu17-Ala21 and Lys28-Val39 form two discontinuous β strands, termed β1 and β2, respectively, upon binding to GM1/DMPC vesicles (**Fig. 2B, Fig. S2, Table S2**). On the other hand, the segment Asp1–Glu11 was not observed, indicating structural disorder and/or high mobility of the N-terminal segment. In general, the β strands of Aβ peptides are stabilized through both intra-or intermolecular hydrogen-bonding interactions. To observe molecular contacts among Aβ peptides in the GM1/DMPC-induced structure, long distance information was acquired by DARR/RAD experiments at various mixing times up to 400 ms (**Fig. S3A**). Intermolecular and intramolecular cross peaks were distinguished based on spectral comparison with uniformly ^13^C- and ^15^N-labeled Aβ diluted with unlabeled Aβ (**Figs. S3B and S3C**), thereby identifying 21 intra- and 20 intermolecular cross peaks (**Tables S3 and S4**). These were used as distance constraints for conformational analyses of Aβ molecules bound to GM1/DMPC vesicles. In terms of intermolecular proximity in DARR/RAD correlation spectra, β1–β1 or β2–β2 interactions were suggested, but we could not determine whether these interactions are arranged in parallel or antiparallel orientation. To determine the arrangements, we used a site-specific Aβ spin label as a source of distance information. Upon addition of a 1:10 molar ratio of unlabeled Aβ-Cys-MTSL (1-oxy-2,2,5,5-tetramethyl-d-pyrroline-3-methyl)methanethiosulfonate) to [^13^C, ^15^N]Aβ, the peaks originating from His13, His14, Gln15, Glu22, Ser26, Asn27, and Ile32 of the isotopically labeled Aβ exhibited significant line broadening due to paramagnetic relaxation enhancement (PRE), indicating that these residues are spatially proximal to the C terminus of another Aβ molecule (**Fig. S4**). If the β2 strands were in parallel, intensity attenuation should be observed only for the peaks originating from amino acid residues proximal to the C terminus. However, since the PREs were not observed at the C terminus, the β2 strands are arranged in an antiparallel manner.

**Fig. 2.**
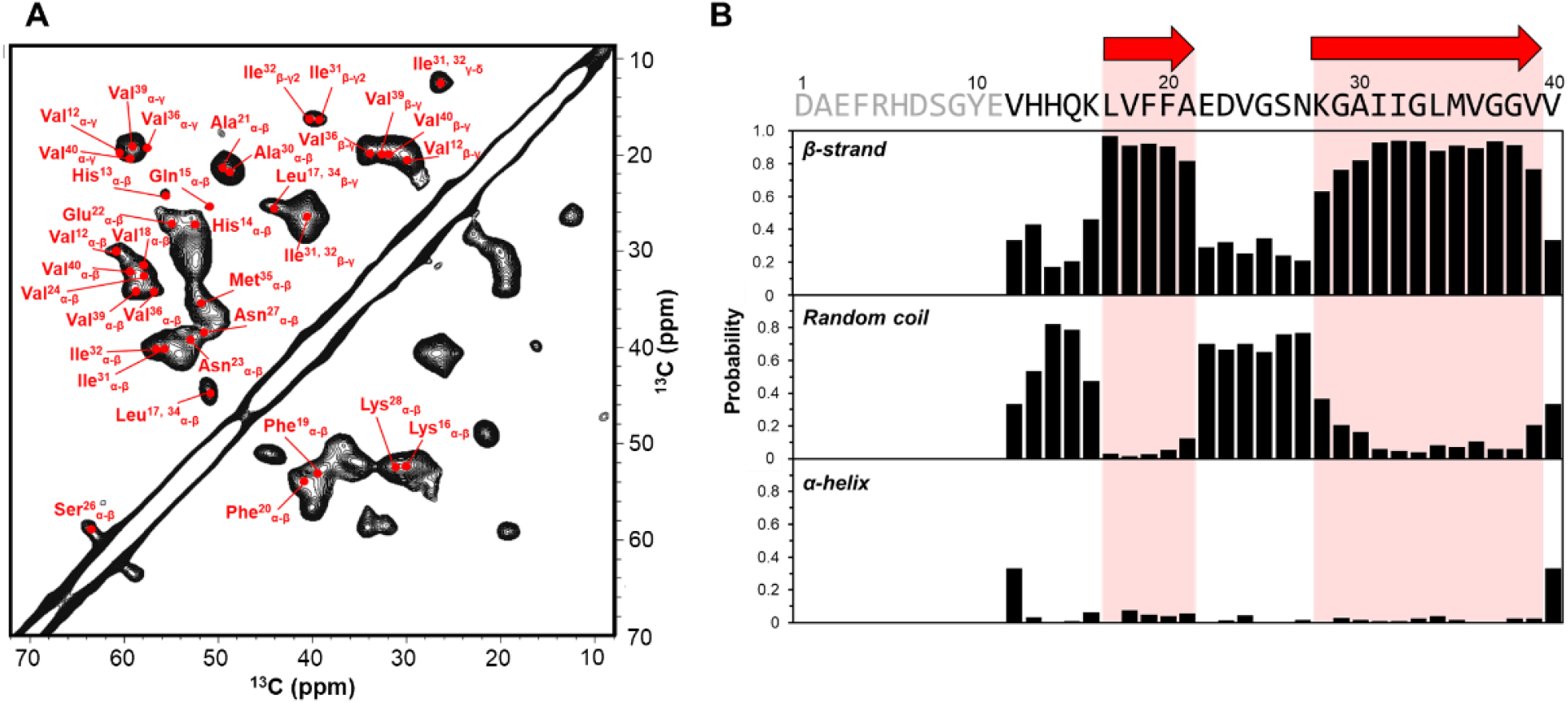
Solid-state NMR characterization of Aβ bound to GM1/DMPC vesicles. (**A**) Aliphatic region of ^13^C–^13^C correlation MAS spectra of [^13^C, ^15^N]Aβ_1–40_ bound to GM1/DMPC vesicles acquired by DARR/RAD with a mixing time of 10 ms. (**B**) Probability of secondary structures (strand, coil, and helix) estimated by TALOS+ analysis according to ^13^C and ^15^N chemical shifts of Aβ_1–40_ bound to GM1/DMPC vesicle. The primary structure of Aβ_1–40_ is presented at the top with red arrows indicating β-strand regions.

As noted above, the Val12–Val40 segment exhibits conformational uniformity among Aβ molecules bound to GM1/DMPC vesicles. Based on the experimentally obtained constraints, we attempted to build a three-dimensional (3D) model of the GM1-boud Aβ assemblage. However, we could not obtain an antiparallel dimer configuration that satisfied all the intermolecular atomic distance constrains obtained from DARR/RAD correlation spectra. Hence, we classified these constraints into two interaction modes and constructed the best compromise model of an Aβ octamer, where eight structurally equivalent Aβ molecules are aligned with antiparallel β1–β1 and β2–β2 interactions (**Figs. S5A**–**C**). This octameric model was used as initial model for MD simulation.

MD calculations were performed using the constraints of dihedral angles and intra- and intermolecular distances estimated based on solid-state NMR data to provide 3D Aβ octamer structures. We calculated the probability of Aβ forming β-structures from the obtained octameric structure, showing that the interaction mode reproduced the β-structure probability calculated from solid-state NMR data using TALOS+ (**Fig. S5D**). The most characteristic feature of the obtained 3D structural model of the Aβ octamer is alternative formation of β1–β1 and β2–β2 hydrogen-bonding interactions, giving rise to two layers of discontinuous antiparallel β-structures (designated as β1- and β2-layers, respectively) (**Fig. 3A**). The final β1 and β2 strands were composed of Lys16-Ala20 and Ile31-Val36, respectively. In β1-layer, intermolecular hydrogen bonds were identified between the NH of Val18 and the CO of Val18, and the CO of Lys16 and the NH of Phe20 (**Fig. 3B**). In β2-layer, intermolecular hydrogen bonds were found between the NH of Gly33 and the CO of Gly33, and between the CO of Ile31 and the NH of Met35 (**Fig. 3C**).

**Fig. 3.**
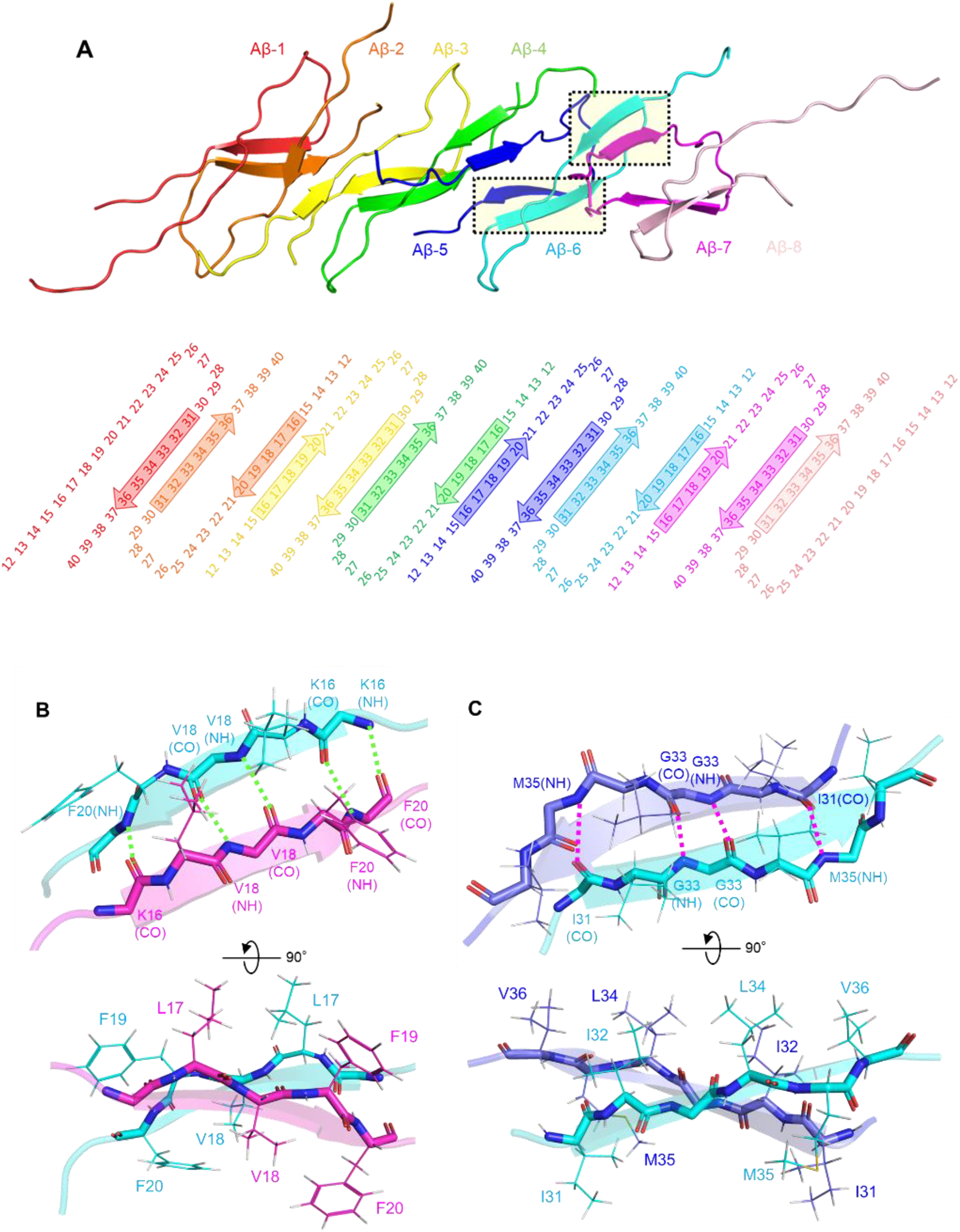
Antiparallel β-structure of Aβ assemblage on GM1/DMPC vesicles. (**A**) 3D structure (upper) and diagrammatic representation (lower) of octameric Aβ_12–40_. Close view of intermolecular β1–β1 (**B**) and β2–β2 (**C**) interactions.

The N-terminal segment (Asp1-Glu11) was not included in the structural model because it is disordered and flexible. We thus examined the possible effect of deletion of N-terminal segments on antibody binding (**Fig. S6**). We confirmed that Aβ_10–40_ lacked reactivity with 6E10. Interestingly, this truncated mutant still retained binding capacity to 4396C when bound to GM1/DMPC vesicles. This indicates that the disordered N-terminal segments are not integral parts of the epitope recognized by 4396C, suggesting that the β1-layer formed on GM1-containing membranes is solvent exposed constituting the conformational epitope to this antibody.

## Discussion

The unique Aβ species tightly associated with GM1 have long been reported to act as an endogenous seed for amyloid fibril formation in the AD brain (*12*). However, it has only been characterized as a conformational epitope recognized by a unique anti-Aβ antibody 4396C (*13*); its structural entity remains largely elusive. The present solid-state NMR study identified a unique antiparallel β-structural assemblage of Aβ specifically formed on GM1 ganglioside clusters. Despite previous structural analyses of amyloid fibrils formed in the presence of lipid membranes (*33, 34*), this study is the first to uncover the structure of nonfibrillar assemblage trapped on GM1-containing membranes.

In contrast to the cross-β structure comprised of in-register parallel β-sheets commonly shared among Aβ fibrils, the assemblage visualized in this study is characterized by a double-layered antiparallel β-structure formed through alternative β1–β1 and β2–β2 hydrogen-bonding interactions along the long axis (**Fig. 3**). It has been reported that a D23N-Aβ_1–40_ mutant can form fibrils characterized by a double-layered in-register highly-stacked antiparallel structure (*35*), structurally distinctive from the loosely-packed GM1-bound Aβ assemblage identified in this study. Aβ peptides corresponding to the β1 strand, such as Aβ_16–22_ and Aβ_11–25_, have been reported to form single-layered in-register antiparallel β-sheets (*36-38*).

Intriguingly, in the antiparallel β-assemblage identified in the present study, the distance between the two layers is almost the same as the dimensions of the GM1 glycan (**Fig. 4**). Antibody-binding data showed that only the β1-layer is antibody accessible, reaching out from the glycan cluster, whereas the β2-layer in inwardly set on the membrane. The previously reported antiparallel fibrils, including the D23N-Aβ_1–40_ fibril, are considered transient intermediate species which eventually evolved into parallel fibrils (*35, 39, 40*). In contrast, the nonfibrillar assemblage on the GM1 membrane appeared stable: our experimental data show that the antiparallel β-assemblage itself was not transformed into ThT-reactive fibrils on GM1 membranes, but could promote conversion of monomeric Aβ into fibrils (**Fig. 1**). The antiparallel β-sheet of Aβ_16–22_ has been reported to catalyze Aβ_1–40_ aggregation through a surface-catalyzed secondary nucleation mechanism (*38*). In mature Aβ fibrils, the hydrophobic surface of cross-β structure appears as pivotal for secondary nucleation (*41-43*). In the double-layered Aβ assemblage identified in the present study, the β1-layer provides a solvent-exposed hydrophobic surface on the GM1-glycan cluster. Based on these data, this β1-layer provides a catalytic surface for Aβ fibril formation.

**Fig. 4.**
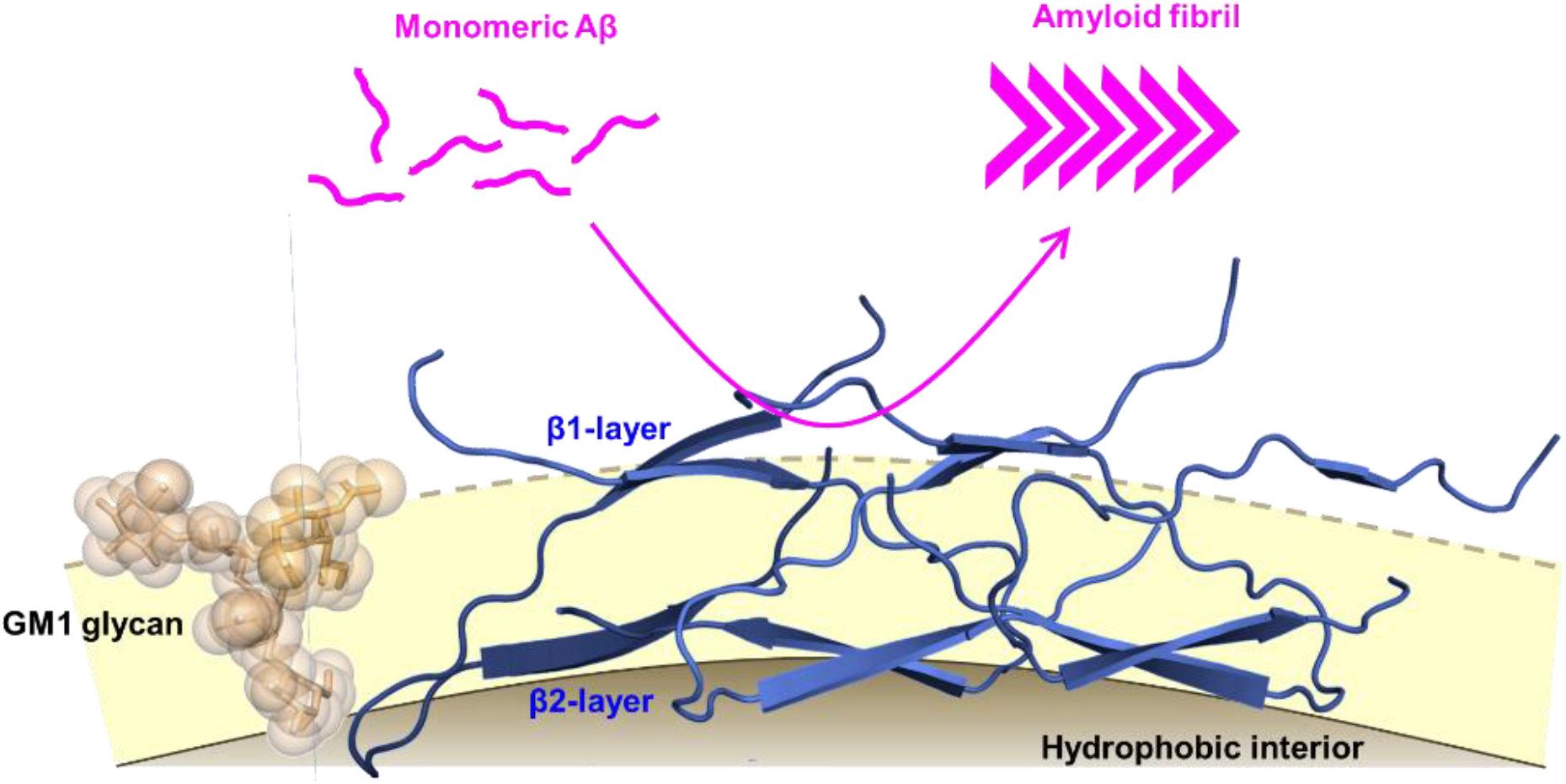
Schematic drawing of Aβ assemblage on GM1-containing membrane which catalytically promotes amyloid fibrillization. Distance between β1- and β2-layers of the assemblage is almost the same as the GM1 glycan dimension. The β1-layer provides a catalytic hydrophobic surface evoking fibril formation in GM1-sugar clusters.

The present data indicate that the hydrophobic surface of β1-layer is exposed, without being covered by the GM1 glycan clusters, and accessible for 4396C binding, suggesting that the conformational epitope recognized by 4396C is located in the antiparallel β1 strands. Noteworthy, aside from GM1-boud-Aβ, 4396C binds an Aβ fibril specifically at its end, thereby preventing fibril elongation in vitro (*13*). This indicates that the Aβ molecule at the fibril end is conformationally distinct from the remaining fibril parts but shares structural similarity with GM1-bound Aβ species in terms of the β1 conformational state. Recently various therapeutic antibodies have been developed for specifically targeting oligomeric or fibrillar Aβ aggregates (*44*). These antibodies are supposed to multivalently bind the N-terminal segments of assembling Aβ molecules. Hence, the 4396C epitope, i.e., the antiparallel β1-structure found in the GM1-bound Aβ assemblage, can be an alternative therapeutic target for suppressing fibril formation in the membrane environment and at the fibril end. Our findings offer structural clues for designing drugs targeting catalytically active conformational species of Aβ for the development of anti-AD therapeutics.

## Materials and Methods

### Preparation of Aβ solution

Expression and purification of uniformly ^13^C- and ^15^N-labeled Aβ_1–40_ was performed as previously described (*22*). The unlabeled Aβ_1–40_ peptide with an extra cysteine residue at its C terminus (Aβ-Cys) was expressed and purified according to the protocol used for wild-type Aβ_1–40_ with slight modifications. The reaction of Aβ-Cys with the nitroxide spin label MTSL (Toronto Research Chemicals) was carried out as previously described (*23*). Synthetic Aβ_1–40_ labeled with ^13^C/^15^N selectively at Val36 or Val40 was purchased from AnyGen Co. N-terminally truncated Aβ mutant, Aβ_10–40_, was purchased from ABclonal Inc. Both recombinant and synthetic Aβ proteins were dissolved at an approximate concentration of 5 mM in 0.1% (v/v) ammonia solution. This Aβ solution was used for preparing samples in subsequent experiments.

### Preparation of small unilamellar vesicle suspension

Powdered DMPC and ganglioside GM1 were purchased from Avanti Polar Lipids Inc. and Carbosynth Ltd., respectively. Small unilamellar vesicles (SUV) composed of GM1 and DMPC were prepared by dissolving 2.8 mg of DMPC and 26.0 mg of GM1 in 1 mL of methanol/chloroform (1:1) solution (molar ratio of DMPC:GM1, 2:8). The solvents were removed from the DMPC/GM1 suspension by nitrogen gas, followed by complete vacuum drying. The dried DMPC/GM1 sample was resuspended in a total of 2 mL of 5 mM potassium phosphate buffer (pH 7.4) and homogenized by six cycles of successive freezing in liquid nitrogen, thawing at 50°C, vortexing at room temperature, and subsequently sonicated for 10 min (2 min, 5 times) using a probe-type sonicator. Metal debris from the probe’s titanium tip was removed by centrifugation. DMPC SUV was prepared in the same manner. The supernatants, i.e., GM1/DMPC suspension (containing 2 mM DMPC and 8 mM GM1) and DMPC suspension (containing 10 mM DMPC), were used for subsequent experiments.

### Preparation of membrane-bound Aβ samples

An aliquot (0.1 mL) of Aβ/ammonium solution was mixed with 1 mL of the DMPC/GM1 suspension and/or the DMPC suspension on ice. After adjusting the pH to 7.4, each mixture was immediately subjected to ultracentrifugation at 604,000 × g for 6 h at 4°C. The supernatant was removed, and the precipitation was immediately lyophilized. A lyophilizate of the Aβ_1–40_/DMPC/GM1 fraction was subjected to solid-state NMR spectroscopy, TEM, ThT assay, and dot blot assay. The lyophilizate of the Aβ_1–40_/DMPC fraction was subjected to ThT assay and dot blot assay; that of Aβ_10–40_/DMPC was subjected to dot blot assay only.

### Solid-state NMR experiments

All solid-state NMR measurements of the lyophilizate of Aβ_1–40_/DMPC/GM1 fraction were carried out under a 14.1-T magnetic field (^1^H resonant frequency of 600 MHz) using a Bruker Avance-600 spectrometer equipped with a ^1^H-^13^C-^15^N triple resonance 2.5-mm magic angle spinning (MAS) probe. The MAS rate was actively controlled at 13.5 kHz ± 5 Hz using a Bruker MAS II automatic controller. Net sample temperature was controlled at 15°C using a variable temperature controller by considering sample heating due to MAS. Representative radio frequency (RF) fields for ^1^H and ^13^C were 100 kHz and 88 kHz, respectively. Samples were packed into 4-mm space at the sample tube center using an original Diflon spacer of 1-mm thickness to maintain RF homogeneity. ^13^C chemical shifts were referenced externally to the ^13^C CH signal of adamantane at 29.5 ppm on the TMS scale, and ^15^N chemical shifts were referenced to the ^15^N signal of NH_4_Cl at 39.25 ppm on the liquid ammonia scale. Recycle delay was 2 s. A ^1^H heteronuclear decoupling during detection period was achieved using small phase incremental alteration-64 (SPINAL-64) (*45*) using a ^1^H RF field of 100 kHz. Initial magnetizations of rare nuclei were enhanced by using ^1^H-X cross polarization (CP) with 20% amplitude sweep of ^1^H spin locking field (*46*) (X = ^13^C and ^15^N). For ^1^H-^13^C CP, an average ^1^H spin locking field of 86.5 kHz, and a ^13^C spin locking field of 73 kHz were used, while for ^1^H-^15^N CP, an average ^1^H spin locking field of 66.5 kHz, and ^15^N spin locking field of 53 kHz were used. Contact times (CTs) were 1,200 and 700 μs for ^13^C and ^15^N, respectively.

The two-dimensional (2D) ^13^C–^13^C correlation was achieved by DARR (*30*)/RAD (*31*) experiments. A constant ^1^H RF field of 13.5 kHz corresponding to the spinning rate was applied during the mixing period. A total of 300, 250, 185, and 185 complex *t*_1_ points were acquired with an increment of 20 μs for mixing times of 10, 100, 200, and 400 ms. For each *t*_1_ point, 1,200–2,480 scans were accumulated. ^1^H decoupling during *t*_1_ period was achieved by SPINAL64.

2D NCO and NCA ^13^C–^15^N correlation experiments were carried out by DCP (*32*) with 78 complex points for *t*_1_ time domain with an increment of 148 μs. For each *t*_1_ point, 2,400 scans were accumulated in both experiments. CTs of 700 and 1,900 μs were used for ^1^H-^15^N and ^15^N-^13^C CPs, respectively. ^15^N-^13^C CP was achieved by a ^13^C spin locking field of 20 kHz and an average ^15^N spin locking field of 33.5 kHz with 5% amplitude sweep. CW decoupling at a ^1^H RF field of 100 kHz was applied during ^15^N-^13^C CP. ^13^C π pulse was applied at the middle of the *t*_1_ period to achieve ^13^C-^15^N heteronuclear *J*-decoupling. During the *t*_1_ period, ^1^H decoupling was achieved by two pulse phase modulation (TPPM) (*47*) with a ^1^H RF field of 100 kHz.

NMR data were processed by TOPSPIN2.1 (Bruker Biospin, Japan). FIDs for 2D ^13^C–^13^C DARR correlation experiments were zero-filled to 4 K points both for *t*_1_ and *t*_2_ time domains, respectively and apodised with a trapezoid window function for both *t*_1_ and *t*_2_ time domains, prior to Fourier transformation (FT). FIDs for 2D ^13^C-^15^N correlation experiments were zero-filled to 4 K and 8 K points for *t*_1_ and *t*_2_ time domains, respectively. FIDs were apodised with trapezoid window functions and gaussian broadenings for the *t*_1_ and *t*_2_ time domains, respectively, prior to FT.

The cross-peaks first observed in DARR spectra at a mixing time of 100 ms were assigned to a short-range distance, <5 Å. While those first observed in the DARR spectra at mixing times of 200 and 400 ms were assigned to a long-range distance, 5.0 ± 2.5 Å.

### ThT assay

Each lyophilizate of the Aβ_1–40_/GM1/DMPC and Aβ_1–40_/DMPC fraction was rehydrated with 1 mL of ultrapure water. The Aβ_1–40_ solution was diluted to 20 μM in 5 mM potassium phosphate buffer (pH 7.4) as monomeric Aβ solution. The rehydrated lyophilizate of the Aβ_1–40_/GM1/DMPC and Aβ_1– 40_/DMPC fractions, the GM1/DMPC suspension, the DMPC suspension, the monomeric Aβ solution, and the monomeric Aβ solution with 1%(v/v) rehydrated lyophilizate of the Aβ_1–40_/GM1/DMPC or Aβ_1–40_/DMPC fraction were supplemented with 40 μM ThT from a 2-mM stock solution. These samples were then pipetted into multiple wells (80 μL per well) of a 96-well half-area, low-binding polyethylene glycol coating plate (Corning 3881) with a clear bottom, followed by incubation at 37°C under quiescent conditions in a plate reader (Infinite 200Pro; TECAN). ThT fluorescence was measured through the bottom of the plate with a 430-nm excitation filter and a 485-nm emission filter. ThT fluorescence was analyzed three times per sample.

### TEM

The rehydrated lyophilizate of Aβ_1–40_/GM1/DMPC fraction was subjected to negative-staining EM. Briefly, specimens stained with 2% uranyl acetate on the grid were observed with an 80 kV electron microscope (JEM-1010; JEOL Inc., Tokyo, Japan). Images were collected at a nominal magnification of 100,000×.

### Dot blot assay

Along with mouse monoclonal antibody 4396C (Immuno-Biological Laboratories Co., Ltd.), mouse monoclonal antibodies 6E10 (COVANCE), 4G8 (COVANCE), and BA27 (FUJIFILM Wako Pure Chemical Corporation), directed against amino acid residues 1–16, 17–24, and the C-terminal region of human Aβ_1–40_, respectively, were used for the dot blot assay.

Aβ_1–40_ and Aβ_10–40_ solutions were diluted at 0.1 mM in 5 mM potassium phosphate buffer (pH 7.4). Aβ fibrils were prepared using 0.1 mM Aβ_1–40_ or Aβ_10–40_ by incubation at 37°C under quiescent conditions. Each lyophilizate of the Aβ_1–40_/GM1/DMPC, Aβ_1–40_/DMPC, and Aβ_10–40_/GM1/DMPC fraction was rehydrated with 1 mL of ultrapure water and then diluted at 0.1 mM in 5 mM potassium phosphate buffer (pH 7.4).

Aβ solutions, Aβ fibrils, and the rehydrated lyophilizates were blotted onto nitrocellulose membranes (BioRad) as described previously (*21*). The blots of theses Aβ samples reacted with 4396C (1:2,500), 6E10 (1:10,000), 4G8 (1:10,000), or BA27 (1:10.000), and subsequently with horseradish peroxidase-conjugated antimouse IgG (Cell Signaling Tec). Bound-enzyme activities were visualized with an enhanced chemiluminescence system (GE Healthcare).

### Molecular dynamics simulation

To determine the intermolecular interaction mode in the Aβ assemblage, we performed an MD simulation of the Val12–Val40 segments of Aβ_1–40_, i.e., Aβ_12–40_, in an explicit water solvent together with the constraints obtained from solid-state NMR experiments. Regarding the β2 strands, the intermolecular distance constraints (listed in **Table S4**) could be classified into two sets: In one set (termed set A), the average of the two residue numbers (residue numbers of Atom 1 and Atom 2 in **Table S4**) is about 33 (as shown in **Fig. S5A**), while in the other set (set B), it is about 36 (as shown in **Fig. S5B**). There is an additional intermolecular distance constraint between Phe20 and Ala21 in the β1 strand. Phe20 is also close to Leu34 and Val36 judging from the intramolecular constraints between β1 and β2 (**Table S3**), suggesting that the region around Gly25–Gly29 forms a loop so that Phe20 and Ala21 are close to the Leu34–Val36 segment in one Aβ molecule. Based on the above considerations, an initial structure of the Aβ assemblage had eight Aβ_12–40_ peptides arranged in antiparallel fashion, assuming that Aβ molecular pairs forming intermolecular hydrogen bonds between odd-numbered residues and those between even-numbered residues satisfy sets A and B, respectively (**Fig. S5C)**.

One Aβ assemblage, consisting of eight Aβ_12–40_ peptides, was placed with eight sodium ions as counterions and 52,537 water molecules in a cubic simulation box with a side length of 118.075 Å. The total number of atoms was 161,123. The N-terminus of each Aβ_12–40_ peptide was capped with an acetyl-group to avoid electrostatic repulsion between the N-termini. The AMBER parm14SB force field (*48*) was used for peptides and counter ions. The TIP3P rigid body model (*49*) was used for water molecules adopting the symplectic (*50*) quaternion scheme for rigid body molecules (*51, 52*). In addition, to include the constraints from the solid-state NMR experiment, the DIANA force field (*53*) was applied: the total potential energy *E* is given by:

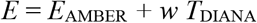

where *E*_AMBER_ is the AMBER potential energy, and *T*_DIANA_ is the DIANA target function. The weight factor *w* for the target function was set as 10 kcal/mol. The electrostatic potential was calculated using the particle mesh Ewald method. The cutoff distance was set as 12 Å for the Lennard–Jones potential.

We used the Generalized–Ensemble Molecular Biophysics program to perform the MD simulation. This program was developed by one of the authors (H. Okumura) and has been used for several protein systems (*54-58*). Reversible multiple time-step MD techniques were also applied (*59*). The time step was *Δt* = 0.5 fs for bonding interactions of peptide atoms, *Δt* = 2.0 fs for nonbonding interactions of peptide atoms, and those between peptide atoms and solvent molecules, and *Δt* = 4.0 fs for the interaction between solvent molecules. Since the symplectic rigid body algorithm was used for water molecules, *Δt* was as long as 4.0 fs (*52*). After energy minimization, an MD simulation in the canonical ensemble was performed for 100 ps at 300 K. The temperature was controlled using the Nosé–Hoover thermostat (*60-62*).

## Acknowledgments

We would like to thank Yukiko Isono (IMS) for help in preparing recombinant proteins. We also thank the Research Equipment Sharing Center at the Nagoya City University, Functional Genomics Facility, NIBB Core Research Facilities, Instrument Center at Institute for Molecular Science, and EM facility in National Institute for Physiological Sciences for technical support.

## Funding

This work was supported in part by JSPS KAKENHI (Grant Numbers JP19K07041 to M.Y.-U., JP21K06040 to S.G.I., JP21K06118 to H.O., JP16K05858 and JP19K05552 to K.N.), by Grant-in-Aid for Research in Nagoya City University Grant Number 2212008 to M.Y.-U. The computation was performed using Research Center for Computational Science, Okazaki Research Facilities, Japan (Projects: 20-IMS-C155, 21-IMS-C172, 22-IMS-C186). Solid-state NMR measurements were conducted in Institute for Molecular Science, supported by Nanotechnology Platform Program <Molecule and Material Synthesis> (JPMXP09S17MS1095, JPMXP09S18MS1055, JPMXP09S18MS1087, JPMXP09S19MS1049) of the Ministry of Education, Culture, Sports, Science and Technology (MEXT), Japan.

## Author contributions

Conceptualization: M.Y.-U., K.Y., K.K., and K.N. designed the research. M.Y.-U. and K.N. prepared the samples. M.Y.-U. and K.N. performed NMR experiments and data analysis. M.Y.-U. performed ThT, EM, and dot blot analyses. S.G.I. and H.O. performed computational analyses. M.Y.-U., H.O., K.K., and K.N. contributed to manuscript writing. All authors contributed to discussions throughout the research and participated in editing or commenting on the manuscript.

## Competing interests

Authors declare that they have no competing interests.

## Data and materials availability

All data needed to evaluate the conclusions in the paper are present in the paper and/or Supplementary Materials. Assigned chemical shift data for Aβ_1–40_ bound to GM1/DMPC vesicle were deposited in the BMRB under the accession number 36495. The atomic coordinates and structure factors are deposited in the Protein Data Bank with accession code 7Y8Q. Additional data related to this paper may be requested from the authors.

## Notes

### Competing Interest Statement

The authors have declared no competing interest.

