## Supplemental Figures and Tables for "The double-layered structure of amyloid-β assemblage on GM1-containing membranes catalytically promotes fibrillization"

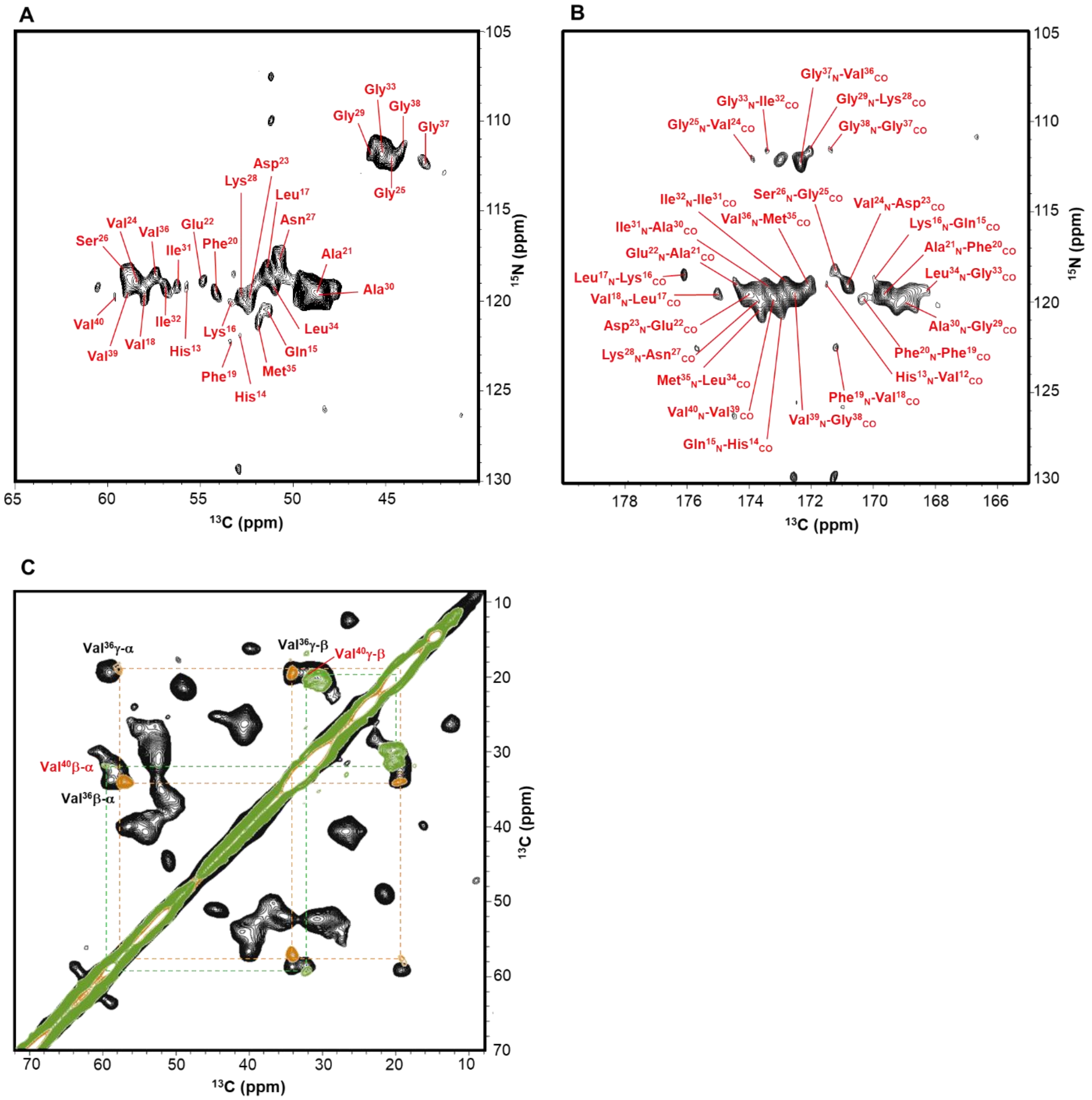


**Fig. S1. NMR assignments of Aβ bound to GM1/DMPC vesicles.** (**A**) 2D NCA and (**B**) 2D NCO $^{13}C$–$^{15}N$ correlation MAS spectra of [$^{13}C$, $^{15}N$]$A\beta_{1–40}$ bound to GM1/DMPC vesicle. (**C**) Aliphatic region of $^{13}C$–$^{13}C$ correlation MAS spectra of [$^{13}C$, $^{15}N$]$A\beta_{1–40}$ (black), [$^{13}C$, $^{15}N$–Val36]$A\beta_{1–40}$ (orange), and [$^{13}C$, $^{15}N$–Val40]$A\beta_{1–40}$ (green) bound to GM1/DMPC vesicle acquired by DARR/RAD with a mixing time of 10 ms.

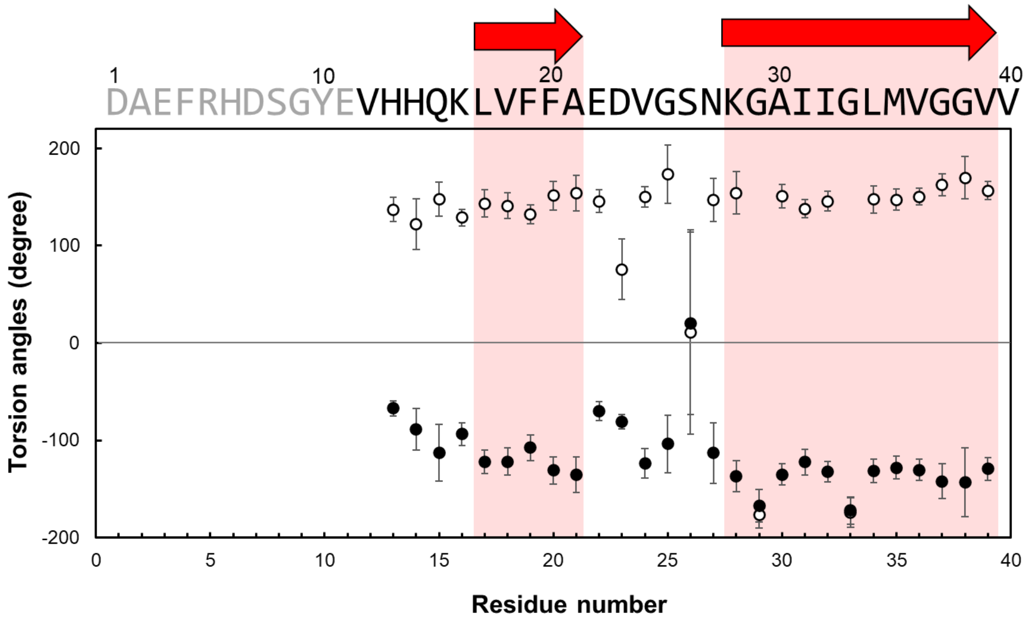


**Fig. S2 Secondary structure of Aβ bound to GM1/DMPC vesicles.** Dihedral angles [φ (open circle), ψ (filled circle)] estimated by TALOS+ analysis according to $^{13}C$ and $^{15}N$ chemical shifts of $Aβ_{1–40}$ bound to GM1/DMPC vesicle. Primary structure of $Aβ_{1–40}$ with red arrows indicating β-strand regions at the top.

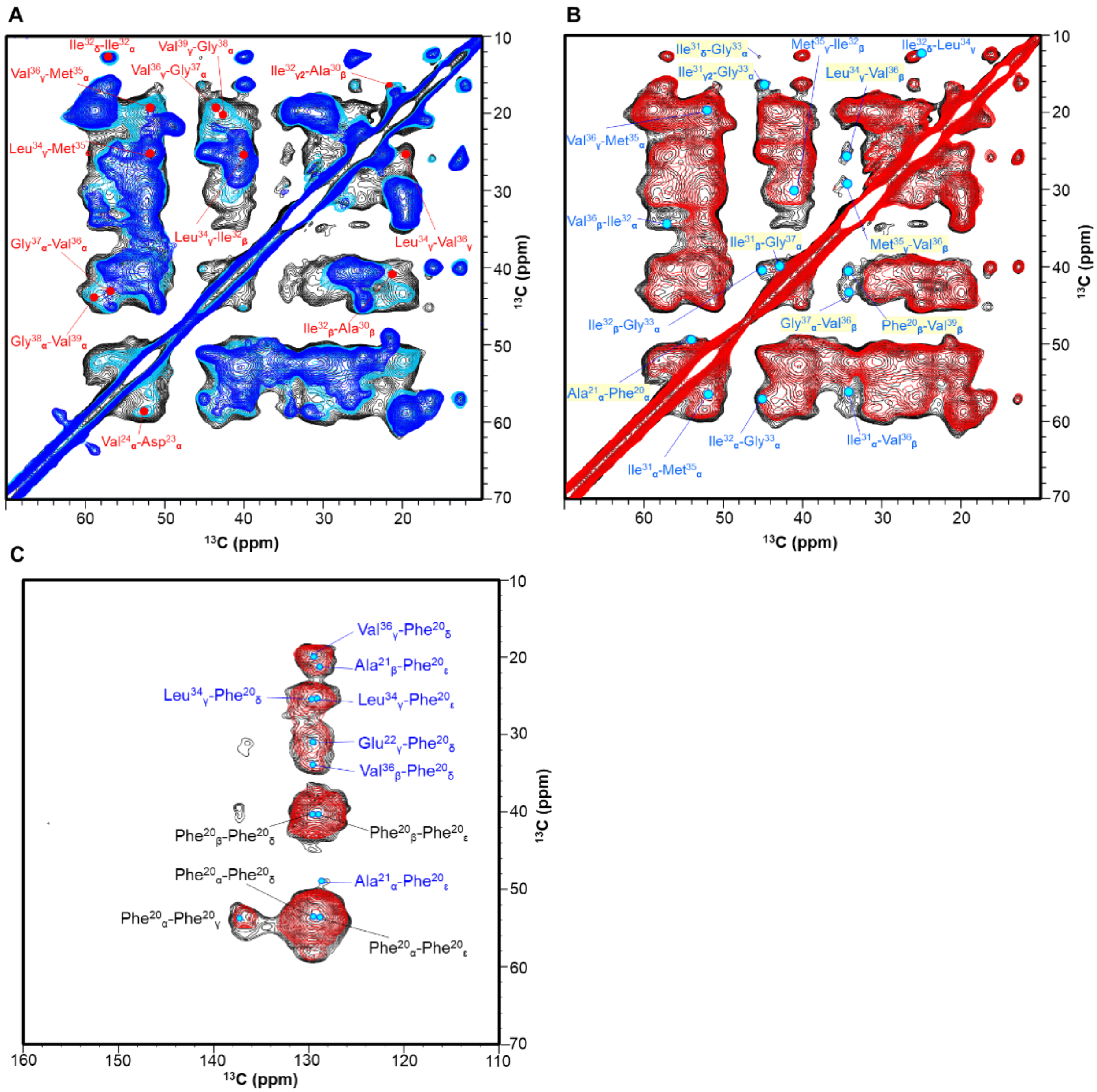


**Fig. S3. Intra- and intermolecular interactions of Aβ bound to GM1/DMPC vesicles.** (**A**) Aliphatic region of $^{13}C$–$^{13}C$ correlation MAS spectra of [$^{13}C$, $^{15}N$]A$\beta_{1\text{–}40}$ bound to GM1/DMPC vesicles acquired by DARR/RAD with mixing times of 100 ms (blue), 200 ms (cyan), and 400 ms (black). (**B**) Aliphatic and (**C**) aromatic regions of $^{13}C$–$^{13}C$ correlation MAS spectra of diluted A$\beta_{1\text{–}40}$ ([$^{13}C$, $^{15}N$]Aβ:unlabeled Aβ = 1:1) bound to GM1/DMPC vesicles acquired by DARR/RAD with a mixing time of 400 ms (red) (4,000 scans were accumulated for each $t_1$ point) superimposed on those of [$^{13}C$, $^{15}N$]A$\beta_{1\text{–}40}$ (black).

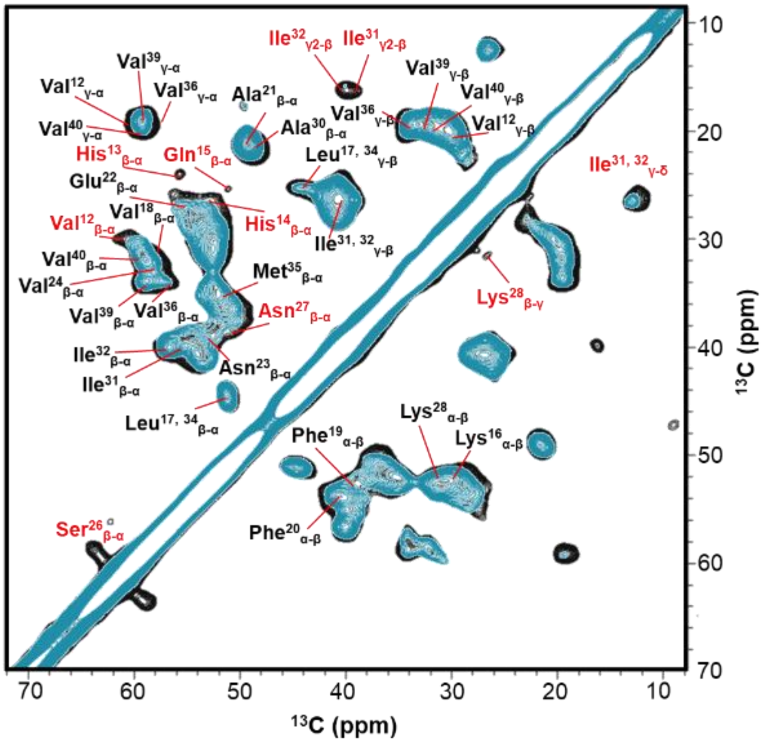


**Fig. S4. Intermolecular interactions of Aβ bound to GM1/DMPC vesicles.** $^{13}C$–$^{13}C$ correlation MAS spectra of [$^{13}C$, $^{15}N$]$A\beta_{1\text{–}40}$ bound to GM1/DMPC vesicles acquired by DARR/RAD with a mixing time of 10 ms in the presence (cyan) and absence (black) of 0.1 molar equivalents of unlabeled Aβ-Cys-MTLS.

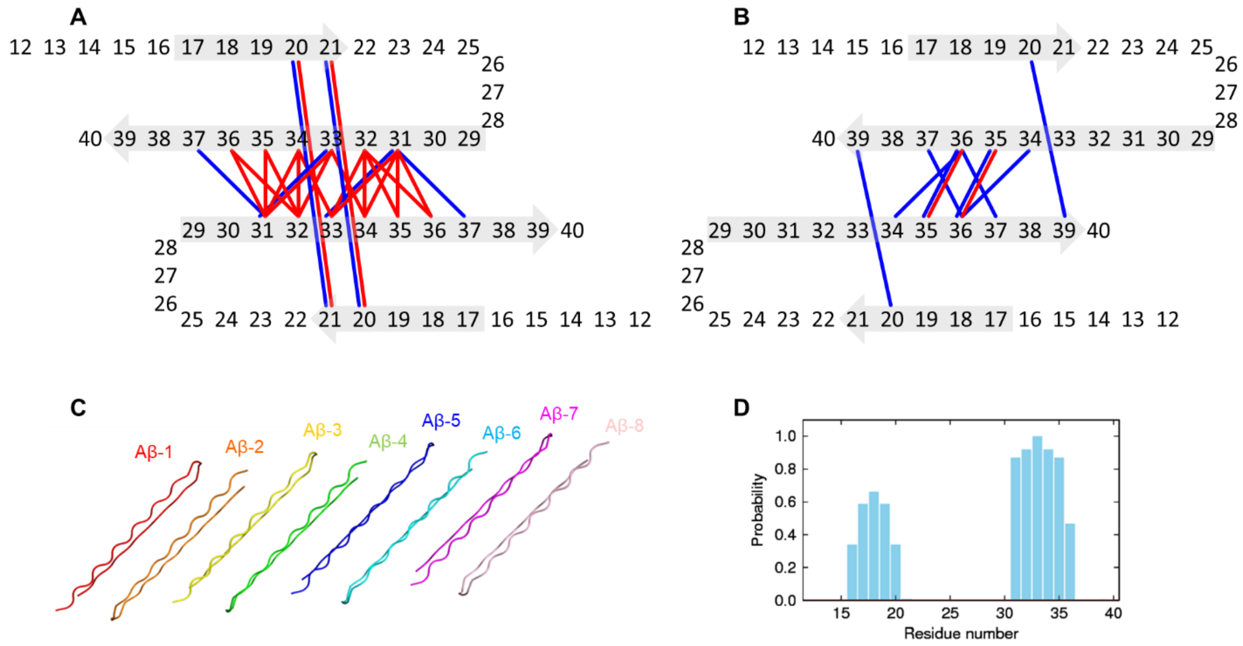


**Fig. S5. MD simulation of Aβ bound to GM1/DMPC vesicles.** (**A** and **B**) Two possible interaction modes in $A\beta_{12\text{–}40}$ dimers based on intermolecular restraints. Blue and red lines indicate long and short distance intermolecular restraints, respectively. (**C**) 3D model of the initial octameric $A\beta_{12\text{–}40}$ structure. (**D**) Probability of β-structures of the octameric Aβ obtained by MD simulation.

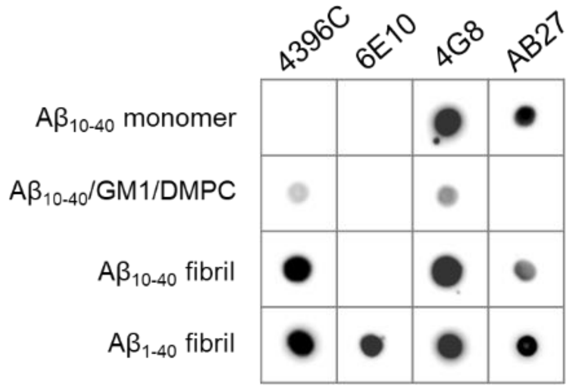


**Fig. S6 Dot blot assay of $A\beta_{10–40}$ with monoclonal anti-Aβ antibodies.** Monomeric $A\beta_{10–40}$, the rehydrated lyophilizate of the $A\beta_{10–40}$/GM1/DMPC fraction, and preprepared $A\beta_{10–40}$ and $A\beta_{1–40}$ fibrils were blotted.

**Table S1. Summary of $^{13}$C, $^{15}$N isotropic chemical shifts with peak assignments for A$\beta_{1-40}$ bound to GM1/DMPC vesicles**

| Residue | Chemical shifts (ppm) | | | | | | | |
|---|---|---|---|---|---|---|---|---|
| | CO | C$\alpha$ | C$\beta$ | C$\gamma$ | C$\delta$ | C$\varepsilon$ | C$\gamma$2 or C$\zeta$ | N |
| V12 | 171.50 | 60.32 | 30.14 | 20.07 | | | | |
| H13 | | 55.71 | 24.00 | 131.76 | 115.73 | | | 119.19 |
| H14 | 173.04 | 52.87 | 26.59 | 131.42 | 117.73 | | | 121.90 |
| Q15 | 169.96 | 51.26 | 25.50 | 33.38 | 173.25 | | | 120.54 |
| K16 | 176.11 | 53.36 | 30.75 | | | | | 120.11 |
| L17 | 174.98 | 51.42 | 44.60 | | | | | 118.38 |
| V18 | 171.18 | 58.05 | | | | | | 119.46 |
| F19 | 170.37 | 53.36 | 39.53 | 137.74 | | | | 122.33 |
| F20 | 169.64 | 54.17 | 40.42 | 137.41 | 130.05 | 128.92 | 127.30 | 119.51 |
| A21 | 174.46 | 49.07 | 21.40 | | | | | 119.46 |
| E22 | 173.72 | 54.98 | 27.11 | 30.87 | 176.08 | | | 118.92 |
| D23 | 170.85 | 52.47 | 38.72 | 176.24 | | | | 119.89 |
| V24 | 173.93 | 58.38 | 32.41 | | | | | 118.92 |
| G25 | 171.18 | 44.67 | | | | | | 112.25 |
| S26 | | 59.03 | 63.60 | | | | | 118.36 |
| N27 | 173.72 | 50.69 | 39.28 | 173.25 | | | | 117.83 |
| K28 | 172.07 | 52.79 | 31.52 | 26.79 | | | | 119.89 |
| G29 | 169.08 | 45.78 | | | | | | 112.13 |
| A30 | 173.40 | 48.43 | 21.40 | | | | | 119.57 |
| I31 | 172.74 | 56.27 | 39.89 | 26.55 | 12.63 | | 16.23 | 119.13 |
| I32 | 173.44 | 56.92 | 40.62 | 26.55 | 12.63 | | 16.23 | 119.24 |
| G33 | 168.35 | 45.19 | | | | | | 111.82 |
| L34 | 173.65 | 51.01 | 44.60 | 25.45 | | | | 119.00 |
| M35 | 172.05 | 51.9 | 35.28 | 29.66 | | | | 121.14 |
| V36 | 172.31 | 57.16 | 34.11 | 19.63 | | | | 118.54 |
| G37 | 171.34 | 42.93 | | | | | | 111.64 |
| G38 | 172.55 | 43.98 | | | | | | 111.17 |
| V39 | 173.20 | 58.82 | 33.95 | 19.47 | | | | 119.46 |
| V40 | 177.80 | 59.43 | 32.09 | 20.12 | | | | 119.78 |

**Table S2. Backbone torsion angles of A$\beta_{1-40}$ bound to GM1/DMPC vesicle predicted by TALOS+**

| Residue | Predicted values (degree) | | | |
|---|---|---|---|---|
| | $\Phi$ | $\Delta\Phi$ | $\psi$ | $\Delta\psi$ |
| H13 | -67.1 | 7.8 | 137.1 | 12.6 |
| H14 | -88.8 | 21.4 | 122.2 | 26.2 |
| Q15 | -112.7 | 29.3 | 147.7 | 17.4 |
| K16 | -93.7 | 11.3 | 128.9 | 8.5 |
| L17 | -122.1 | 11.8 | 143.5 | 13.9 |
| V18 | -122.0 | 14.1 | 141.1 | 13.4 |
| F19 | -107.5 | 13.3 | 132.4 | 9.7 |
| F20 | -130.9 | 13.9 | 151.3 | 14.7 |
| A21 | -135.4 | 18.6 | 154.3 | 18.2 |
| E22 | -70.1 | 10.1 | 145.8 | 11.9 |
| D23 | -81.0 | 7.2 | 75.6 | 31.2 |
| V24 | -123.9 | 15.4 | 150.3 | 10.5 |
| G25 | -103.8 | 29.3 | 173.5 | 30.3 |
| S26 | 20.2 | 94.1 | 11.1 | 105.1 |
| N27 | -113.2 | 31.4 | 146.9 | 22.2 |
| K28 | -137.1 | 15.9 | 154.3 | 21.9 |
| G29 | -167.1 | 17.0 | -176.8 | 13.2 |
| A30 | -135.0 | 11.2 | 150.7 | 11.9 |
| I31 | -122.4 | 13.4 | 137.9 | 9.2 |
| I32 | -132.4 | 10.6 | 145.6 | 10.2 |
| G33 | -172.2 | 14.1 | -174.2 | 15.4 |
| L34 | -131.8 | 12.1 | 147.5 | 14.3 |
| M35 | -128.1 | 11.9 | 147.3 | 10.9 |
| V36 | -130.7 | 10.9 | 150.4 | 8.3 |
| G37 | -142.0 | 18.0 | 162.5 | 11.5 |
| G38 | -143.3 | 35.5 | 169.9 | 21.9 |
| V39 | -129.5 | 11.8 | 156.5 | 9.4 |

**Table S3. Intramolecular restraints used for structure calculations of A$\beta_{1-40}$ bound to GM1/DMPC vesicles**

Short distance

| Atom 1 | Atom 2 | distance |
|---|---|---|
| I32Cδ | I32Cα | < 5.0 Å |
| D23Cα | V24Cα | < 5.0 Å |
| V36Cα | G37Cα | < 5.0 Å |
| V39Cγ | G38Cα | < 5.0 Å |
| V39Cα | G38Cα | < 5.0 Å |
| A30Cβ | I32Cβ | < 5.0 Å |
| E22Cγ | F20Cδ | < 5.0 Å |
| L34Cγ | F20Cε | < 5.0 Å |
| V36Cγ | F20Cδ | < 5.0 Å |
| L34Cγ | F20Cδ | < 5.0 Å |
| V36Cβ | F20Cδ | < 5.0 Å |
| I31CO | I31Cγ2 | < 5.0 Å |
| I31CO | I32Cβ | < 5.0 Å |
| V36CO | V36Cγ | < 5.0 Å |

Long distance

| Atom 1 | Atom 2 | distance |
|---|---|---|
| A30Cβ | I32Cβ | 5.0 ± 2.5 Å |
| L34Cγ | I32Cβ | 5.0 ± 2.5 Å |
| L34Cγ | V36Cγ | 5.0 ± 2.5 Å |
| L34Cγ | M35Cα | 5.0 ± 2.5 Å |
| V36Cγ | M35Cα | 5.0 ± 2.5 Å |
| V36Cγ | G37Cα | 5.0 ± 2.5 Å |
| I32Cγ2 | A30Cβ | 5.0 ± 2.5 Å |

**Table S4. Intermolecular restraints used for structure calculations of A$\beta_{1\text{–}40}$ bound to GM1/DMPC vesicles**

Short distance

| Atom 1 | Atom 2 | distance |
|---|---|---|
| M35Cα | V36Cγ | < 5.0 Å |
| I31Cα | M35Cα | < 5.0 Å |
| I31Cα | V36Cβ | < 5.0 Å |
| I31CO | L34Cγ | < 5.0 Å |
| I32Cα | V36Cβ | < 5.0 Å |
| I32Cβ | M35Cγ | < 5.0 Å |
| I32Cδ | L34Cγ | < 5.0 Å |
| G33Cα | I32Cα | < 5.0 Å |
| G33Cα | I32Cβ | < 5.0 Å |
| G33CO | I31Cγ2 | < 5.0 Å |
| G33CO | L34Cγ | < 5.0 Å |
| A21Cβ | F20Cε | < 5.0 Å |

Long distance

| Atom 1 | Atom 2 | distance |
|---|---|---|
| G33Cα | I31Cγ2 | 5.0 ± 2.5 Å |
| G33Cα | I31Cδ | 5.0 ± 2.5 Å |
| G37Cα | I31Cβ | 5.0 ± 2.5 Å |
| V36Cβ | L34Cγ | 5.0 ± 2.5 Å |
| V36Cβ | G37Cα | 5.0 ± 2.5 Å |
| V36Cβ | M35Cγ | 5.0 ± 2.5 Å |
| V39Cβ | F20Cβ | 5.0 ± 2.5 Å |
| A21Cα | F20Cα | 5.0 ± 2.5 Å |
